# Quantitative microbiome profiling in lumenal and tissue samples with broad coverage and dynamic range via a single-step 16S rRNA gene DNA copy quantification and amplicon barcoding

**DOI:** 10.1101/2020.01.22.914705

**Authors:** Said R. Bogatyrev, Rustem F. Ismagilov

## Abstract

Current methods for detecting, accurately quantifying, and profiling complex microbial communities based on the microbial 16S rRNA marker genes are limited by a number of factors, including inconsistent extraction of microbial nucleic acids, amplification interference from contaminants and host DNA, different coverage of PCR primers utilized for quantification and sequencing, and potentially biases in PCR amplification rates among microbial taxa during amplicon barcoding. Here, we describe a single-step method that enables the quantification of microbial 16S rRNA gene DNA copies with wide dynamic range and broad microbial diversity, and simultaneous amplicon barcoding for quantitative 16S rRNA gene amplicon profiling of microbiota. The method is suitable for a variety of sample types and is robust in samples with low microbial abundance, including samples containing high levels of host mammalian DNA, as is common in human clinical samples. We demonstrate that our modification to the Earth Microbiome Project (EMP) V4 16S rRNA gene primers expands their microbial coverage while dramatically reducing non-specific mammalian mitochondrial DNA amplification, thus achieving wide dynamic range in microbial quantification and broad coverage for capturing high microbial diversity in samples with or without high host DNA background. The approach relies only on broadly available hardware (real-time PCR instruments) and standard reagents utilized for conventional 16S rRNA gene amplicon library preparation both of which make it amenable for immediate and widespread adoption. Simultaneous 16S rRNA gene DNA copy quantification and amplicon barcoding for multiplexed next-generation sequencing from the same analyzed sample, performed in a combined workflow, reduces the amount of sample needed and reduces time and reagent costs. Additionally, we demonstrate that using our modified 16S rRNA gene primers in a digital PCR (dPCR) format enables precise and exact microbial quantification in samples with very high host DNA background levels without the need for quantification standards. Potential future applications of this approach include: (1) quantitative microbiome profiling in human and animal microbiome research; (2) detection of monoinfections and profiling of polymicrobial infections in tissues, stool, and bodily fluids in human and veterinary medicine; (3) environmental sample analyses (e.g., soil and water); and (4) broad-coverage detection of microbial food contamination in products high in mammalian DNA, such as meat products. We predict that utilization of this approach primarily for quantitative microbiome profiling will be invaluable to microbiome studies, which have historically been limited to analysis of relative abundances of microbes.

## INTRODUCTION

Microbiome analysis has emerged as a prominent research field to improve our understanding of the host-microbiota interactions linked to human disease. Utilization of high-throughput next generation sequencing (NGS) technology in combination with microbial marker gene sequencing (e.g., microbial 16S rRNA gene) has enabled high-diversity and high-depth compositional analyses of microbiomes. NGS-based compositional analyses (relative abundances of the microbiome elements) have dominated the field since their emergence. The limitations of compositional analyses have been gaining broader acknowledgement in the field and a number of quantitative microbiome profiling approaches have been proposed as promising tools for solving the shortcomings of purely compositional analyses. Current quantitative analysis approaches have important limitations: (i) high levels of host DNA interfere with the amplification of target microbial sequences, (ii) coverage of microbial taxa is limited, and (iii) relative quantification cannot provide a complete picture of changes in microbial taxa.

Here, to address the aforementioned limitations of current quantitative analysis methods, we describe an approach that allows simultaneous (with a single aliquot of DNA for each sample) determination of the absolute 16S rRNA gene DNA copy loads with broad dynamic range and enables wide-diversity microbiome profiling in a simplified and broadly-adoptable workflow (Fig. 1). The proposed approach for quantitative 16S rRNA gene amplicon profiling is based on the combination of absolute 16S rRNA gene DNA copy quantification and 16S rRNA gene amplicon sequencing utilizing a real-time PCR amplification readout and amplicon barcoding for NGS performed for the variable V4 region of the prokaryotic 16S rRNA gene sequence amplicon.

**Fig. 1.**
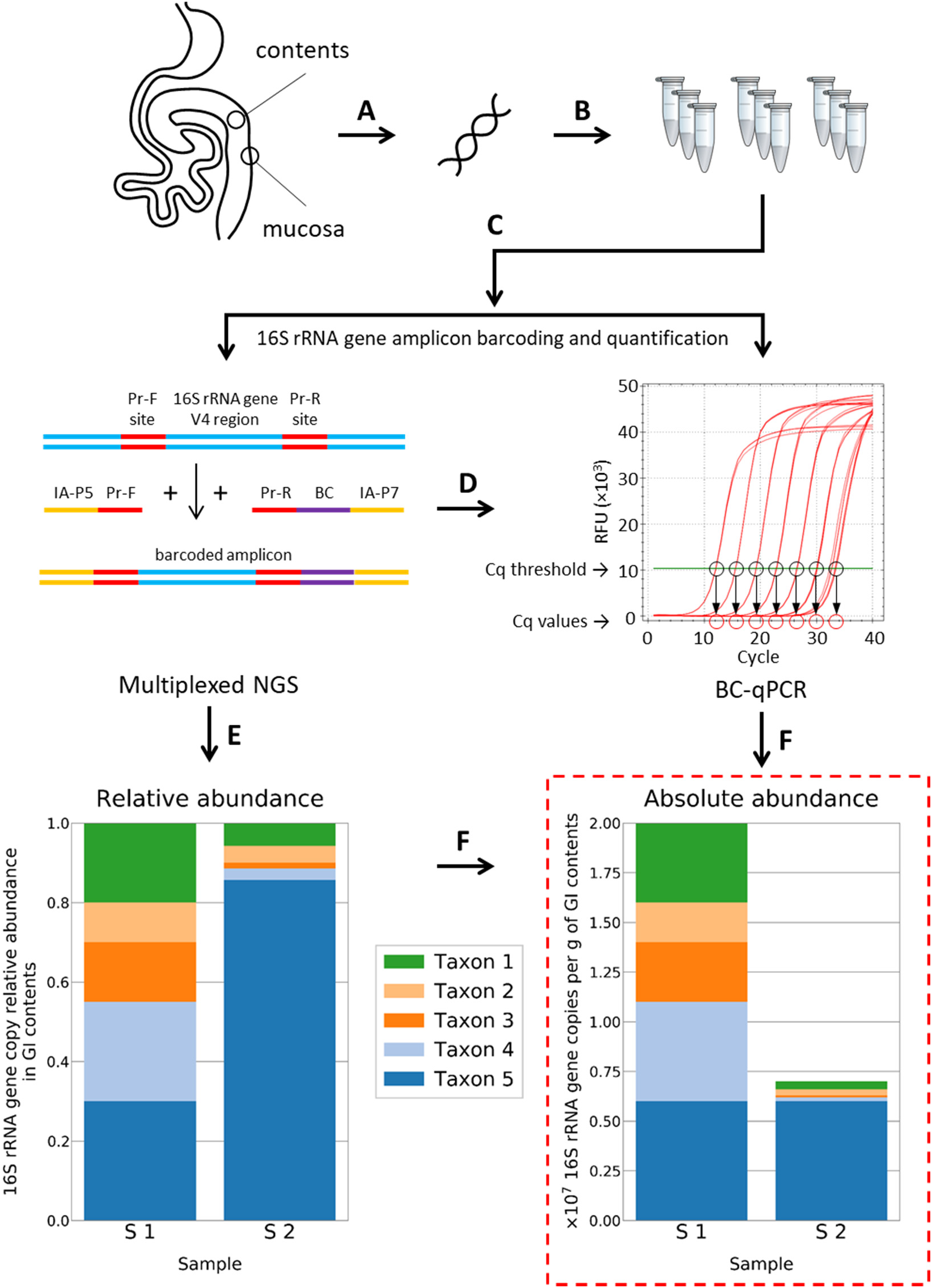
Schematic of the single-step 16S rRNA gene DNA quantification and amplicon barcoding workflow (BC-qPCR) implementation for quantitative microbiome profiling. (**A**) Sample collection and DNA extraction. (**B**) BC-qPCR reactions are prepared in replicates for more accurate quantification and uniform amplicon barcoding. (**C**) Amplification and barcoding of the V4 region of microbial 16S rRNA gene are performed under real-time fluorescence measurements on a real-time PCR instrument. Pr-F – forward primer, Pr-R – reverse primer, IA-P5 and IA-P7 – Illumina adapters P5 and P7 respectively, BC – barcode. (**D**) Quantitative PCR data (Cq values) are recorded. Mock data are shown for illustration. (**E**) Barcoded samples are quantified, pooled, purified, and sequenced on an NGS instrument. (**F**) NGS sequencing results provide data on relative abundances of microbial taxa (mock chart data were constructed only for illustrative purposes). (**G**) Microbial taxa relative abundance profiles are converted to microbial absolute or absolute fold-difference abundance profiles using the absolute or absolute fold-difference data (16S rRNA gene DNA loads) measured in the corresponding samples in step (D) (mock chart data were constructed only for illustrative purposes).

This “single-step” approach is optimized for use in samples with high or low levels of mammalian (e.g., mouse) host DNA and can enable quantitative 16S rRNA gene amplicon profiling of clinical samples, such as stool, gastrointestinal contents or lavage fluid, and mucosal biopsies.

### “Single-step” quantification and library preparation workflow

“Single-step” approach includes the following workflow:

1. Total DNA is extracted and purified from samples (e.g., feces, gastrointestinal contents or aspirates, intestinal mucosa biopsy) using commercially-available kits (Fig. 1A) validated for uniform DNA extraction from complex microbiota (e.g., ZymoBIOMICS) and for quantitative recovery of microbial DNA from samples with microbial loads across multiple orders or magnitude (Fig. S1).
2. PCR reactions are set up with the improved and optimized universal 16S rRNA gene primers (discussed in detail below) containing barcodes and Illumina adapters (Fig. 1B) and conventional commercial reagents for 16S rRNA gene amplicon library preparation. Reactions can be run in replicates to improve the real-time PCR quantification precision and resolution and amplicon barcoding uniformity [1].
3. Amplification and barcoding of the V4 region of the microbial 16S rRNA gene DNA are performed under real-time fluorescence measurements on a real-time PCR instrument (Fig. 1C). We define this approach as “barcoding qPCR” or “BC-qPCR”. Real-time fluorescence monitoring enables terminating the amplification of each sample upon reaching the mid-exponential phase to maximize the amplicon yield and minimize the over-amplification artifacts [2].
4. Quantitative real-time PCR data (Cq values) are recorded (Fig. 1D) and used to calculate the absolute concentration of the 16S rRNA gene DNA copies in each sample (based on the 16S rRNA gene copy standards included within the same BC-qPCR run) or to calculate the absolute fold-differences in the 16S rRNA gene DNA copy load among the samples (in the absence of the standards). These data are further used to calculate the absolute abundances or fold differences in the absolute abundances of individual microbial taxa in the analyzed samples.
5. Barcoded 16S rRNA gene DNA amplicon samples are quantified, pooled, purified, and sequenced on an NGS instrument. NGS sequencing results provide the sequence read and count data which enable taxa identification and calculation of the corresponding taxa relative abundance profiles for the analyzed samples (Fig. 1E).
6. Microbiota relative abundance profiles (from step “E”) are converted to microbiota absolute or absolute abundance fold-difference profiles using the absolute or absolute fold-difference data on 16S rRNA gene DNA loads in the corresponding samples (as measured in the step “D”) (Fig. 1F).

### Anchoring approaches for quantitative sequencing of microbiota

Several different anchoring approaches can be utilized to convert the Cq values obtained from the BC-qPCR assay to total 16S rRNA gene loads in the analyzed samples:

1. Single 16S rRNA gene DNA standard with known template concentration can be included with a 96-well (96 PCR tube) run (Fig. 2A). Alternatively, a single uncharacterized sample from the batch can be analyzed using the companion dPCR assay (discussed below), and thus can serve as a single “standard” anchoring sample. In this scenario all calculations of the absolute concentrations of the remaining samples can be done using the equations 1.1. and 1.2 and would rely on the empirical BC-qPCR efficiency of 95.0% (discussed below), which may or may not hold for any given batch of samples and reagents. Absolute abundance values of total 16S rRNA gene copies can then be converted to the absolute abundances of 16S rRNA gene copies for each individual taxon based on their relative abundance values (obtained from NGS).
2. Two or more 16S rRNA gene DNA standards (serial dilutions) with known template concentrations can be utilized in a similar manner (Fig. 2B). Including more than one standard will allow estimating the exact BC-qPCR efficiency (according to the equations 2.2 and 2.5 or 2.3, 2.4, and 2.5) for any given batch of samples and reagents. Similarly to the first scenario, multiple uncharacterized samples (ideally with distant Cq values) can be quantified using the dPCR assay and used as anchoring standards for the batch of samples. Any of the multiple anchoring points then can be used to calculate the 16S rRNA gene copy load in the remaining samples from the batch using equations 2.1 and 2.6. The conversion of taxa 16S rRNA gene relative abundance values to their absolute abundance values is then performed as in 1.
3. In the absence of absolute anchoring (e.g., quantitative standards or dPCR are not available) the BC-qPCR approach can still provide a valuable information about the fold difference in the absolute load of 16S rRNA gene copies among compared samples (in a single batch and potentially across multiple batches). Such fold difference between samples can be calculated using the equations 3.1 and 3.2 and assuming the empirical BC-qPCR efficiency of 95.0%. Such absolute fold difference values can then be converted to the absolute fold differences among samples for each individual taxon based on their relative abundance values (obtained from NGS).

**Fig. 2.**
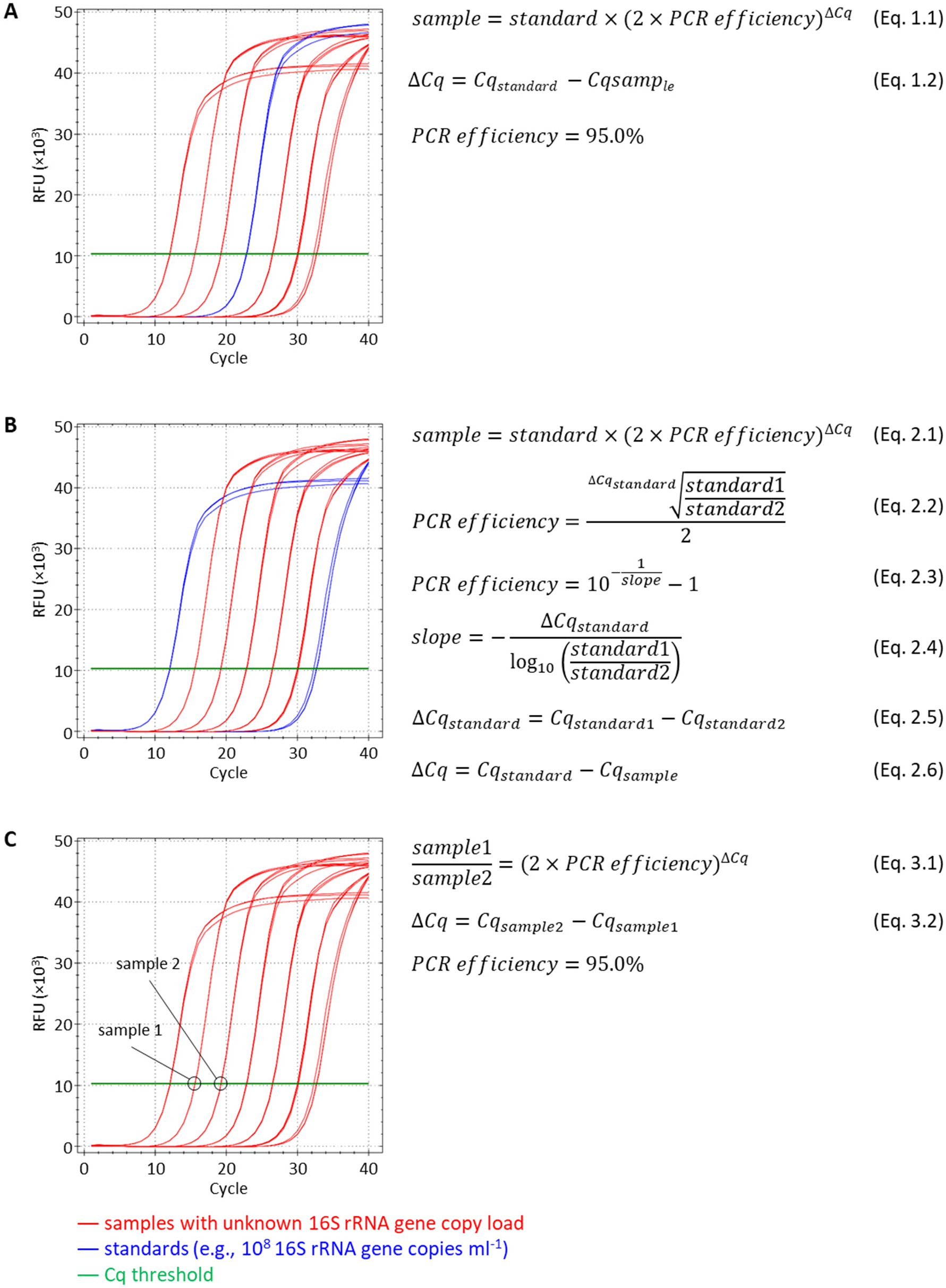
A schematic drawing describing anchoring approaches for deriving the absolute abundances or absolute abundance fold differences implemented with the single-step 16S rRNA gene DNA quantification and amplicon barcoding workflow (BC-qPCR). (A) Anchoring with a single standard and assumed BC-qPCR efficiency. (B) Anchoring with two or more standards and calculated batch-specific BC-qPCR efficiency. (C) Estimation of the absolute fold difference among samples with unknown total 16S rRNA gene DNA copy load in the absence of standards.

To achieve the desired broad dynamic range and coverage of the quantitative 16S rRNA gene amplicon profiling and its robust performance in samples with high or low host DNA background, we: (I) modified the broadly-adopted Earth Microbiome Project (EMP) universal 16S rRNA gene primers for gene-copy quantification in qPCR and dPCR assays and amplicon barcoding in BC-qPCR with high specificity against host DNA; (II) optimized the BC-qPCR parameters to minimize primer dimer formation and host DNA amplification while reducing amplification biases and ensuring uniform amplification of diverse 16S rRNA gene sequences from complex microbiomes; (III) validated the accuracy of the quantitative 16S rRNA gene amplicon profiling obtained using the single-step BC-qPCR approach compared with the quantitative 16S rRNA gene amplicon profiling results obtained using real-time and digital PCR.

## RESULTS

### Optimized primers improve broad-coverage 16S rRNA gene DNA quantification via real-time and digital PCR in the presence of high host DNA background

We first aimed to adapt the Earth Microbiome Project (EMP) 16S rRNA gene amplicon profiling protocol [1, 3] for quantitative microbiota profiling. This protocol is well-known for having broad microbial coverage and has been widely adopted in the field of basic and clinical microbiome research. We hypothesized that by redesigning the EMP forward primer (designated by us as UN00F0) at its 5′ end to start at position 519 (UN00F2) of the V4 region of microbial 16S rRNA gene sequence (Fig. 3A) we would either reduce or eliminate its nonspecific annealing to the mouse and human mitochondrial 12S rRNA gene DNA – the main competing template of mammalian origin identified by amplicon sequencing of PCR products obtained with mouse germ-free tissue DNA. Such change would increase the primer’s specificity for low copy number microbial templates in samples with high content of mouse or human host DNA background. We confirmed the effectiveness of these design considerations by performing qPCR reactions in complex mouse microbiota DNA samples analyzed as-obtained or spiked with GF mouse small-intestine mucosal DNA at 100 ng/μL. The ~200-bp mithochondrial 12S rRNA gene amplicons were absent in the PCR reactions containing high amounts of mouse DNA and using the modified forward primer UN00F2 (Fig. 3B).

**Fig. 3.**
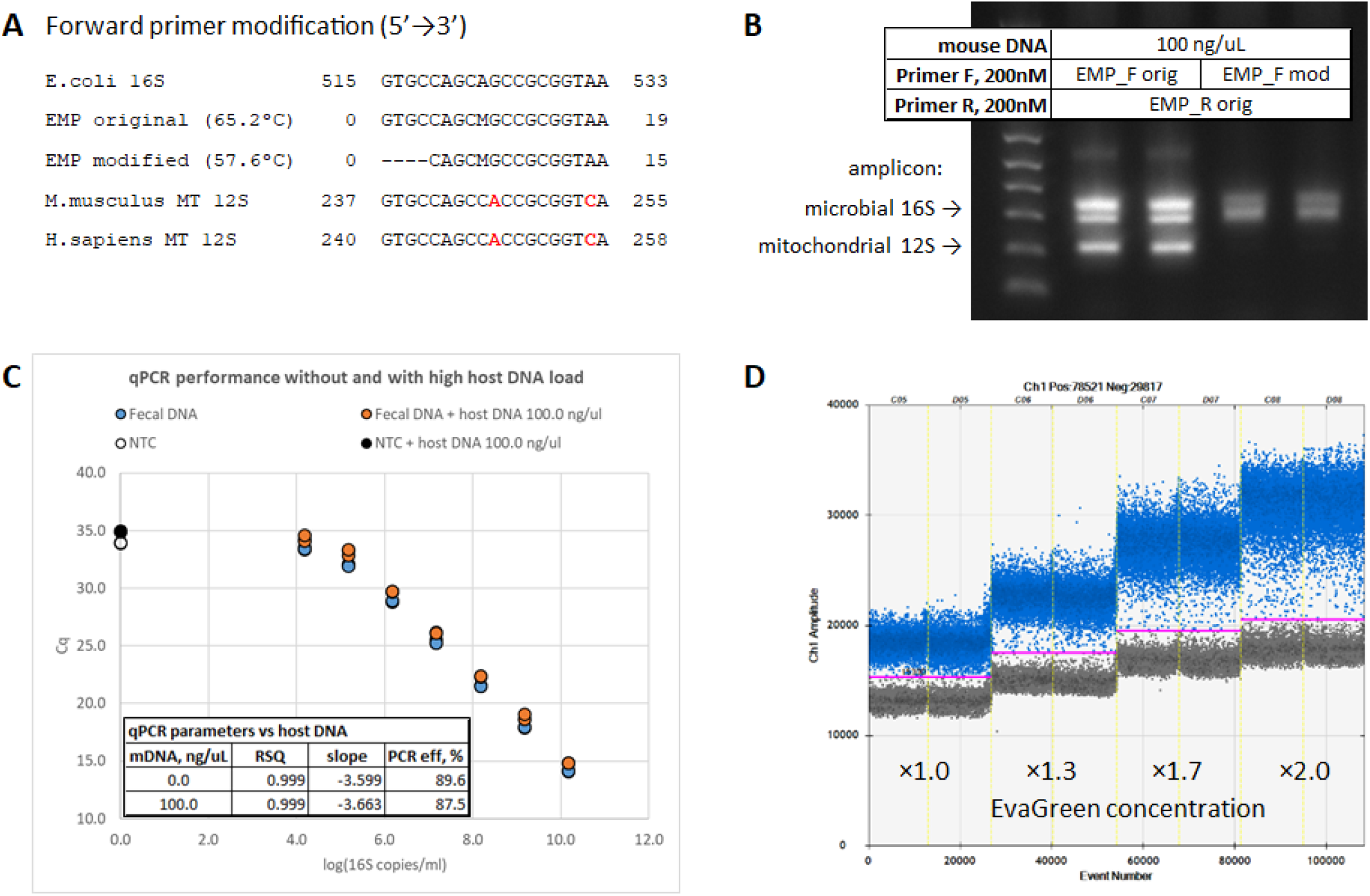
Optimization of the protocol for microbial 16S rRNA gene DNA copy quantification in samples without and with high mammalian DNA background. **(A)** Sequence alignment of the original EMP and modified forward primers targeting the V4 region of microbial 16S rRNA gene are shown with the *E. coli* 16S rRNA gene and mouse and human mitochondrial 12S rRNA gene sequences. **(B)** Amplification products of the complex microbiota DNA sample containing 100 ng/μL of GF mouse DNA obtained with the original EMP or modified forward primers. **(C)** Quantitative PCR reaction performance with the serial 10-fold dilutions of the complex microbiota DNA sample with and without 100 ng/µL of mouse DNA. **(D)** Improvement of the 16S rRNA gene DNA copy ddPCR quantification assay performance in the presence of 100 ng/µL of mouse DNA background as a result of the supplementation of intercalating dye to the commercial droplet digital PCR (ddPCR) master mix.

The efficiency of the quantitative PCR reactions set up with the modified forward primer UN00F2 was similar (and high) with and without the presence of 100 ng/µL of mouse DNA in the template sample (Fig. 3C) demonstrating the robust assay performance.

Our qPCR experiments also suggested that the PCR reactions with high host DNA background are intercalating dye-limited: the increase in total fluorescence (∆-RFU) in each reaction at the end of amplification was lower in samples containing 100 ng/µL of background mouse DNA whereas the total fluorescence levels were similar between samples with and without the background mouse DNA. By combining the use of the new forward primer UN00F2 with the supplementation of commercial reaction mix with additional amounts of intercalating EvaGreen dye improved the digital PCR performance by increasing the separation between negative and positive droplets in the droplet digital PCR (ddPCR) reactions used for quantifying 16S rRNA gene DNA copies in samples with high host DNA background (100 ng/μL) (Fig. 3D). This assay was used to establish or confirm the exact 16S rRNA gene DNA copy numbers in the standard samples, which were further utilized to build the standard curves in the qPCR assays.

Additionally, the modification of the primer set UN00F2 + UN00R0 broadened its taxonomical coverage of the microbial diversity (86.0% Archaea, 87.0% Bacteria) compared with the original EMP primer set UN00F0 + UN00R0 (52.0% Archaea, 87.0% Bacteria) based on the SILVA 16S rRNA gene sequence reference database [4, 5].

### Modified barcoded primers and optimized workflow enable simultaneous 16S rRNA gene DNA copy quantification and amplicon barcoding in samples with high host DNA background

We next aimed to evaluate whether the barcoded UN00F2 + UN00R0 primer set would allow the amplification and amplicon barcoding of specific microbial 16S rRNA gene DNA template in the presence of high host DNA background. It is important to note the two essential—and contrasting—design principles in the BC-qPCR reaction optimization that guided our work:

1. The amplification and barcoding reaction utilizing degenerate 16S rRNA gene primers (whether using the original EMP primers or improved EMP primers) should be conducted at the lowest possible annealing temperature (within the range of annealing temperatures for the specific primer variants within the degenerate primer mixture) to maximize the uniformity of amplification of diverse 16S rRNA gene DNA sequences and eliminate the amplification biases.
2. The amplification and barcoding reaction should be conducted at the highest possible annealing temperature to minimize the primer dimer formation and non-specific host mitochondrial DNA amplification both of which would be competing with specific microbial 16S rRNA gene DNA template for reaction resources (dNTPs, primers, polymerase, intercalating dye). Such competing reactions would inevitably have pronounced effects on the samples containing very low levels of specific microbial template and requiring high numbers of amplification cycles.

Compared with the improved primer set (UN00F2 + UN00R0), the original EMP primer set (UN00F0 + UN00R0) requires a higher annealing temperature to reduce primer dimer formation and amplification of mouse mitochondrial (MT) DNA. Long “overhangs” (carrying the linker and Illumina adapter sequences) at the 5′ end of the forward primer and non-complimentary to the specific 16S rRNA gene DNA template were not sufficient to prevent the EMP primer set from amplifying the mouse MT DNA. At 53.9 °C both primer dimers and MT DNA amplification persisted in the reactions using the EMP primers, which suggested that this primer set would require even higher annealing temperatures (>53.9 °C) to eliminate the amplification artifacts. This in turn will likely introduce amplification biases across a range of specific 16S rRNA gene DNA templates. Using the improved primer set eliminated both artifacts in the reactions conducted at 53.9 °C (Fig. 4A), while some primer dimer formation was still present in the reactions conducted at 52 °C. Thus, the temperature of 54 °C was selected as optimal for the BC-qPCR reaction.

**Fig. 4.**
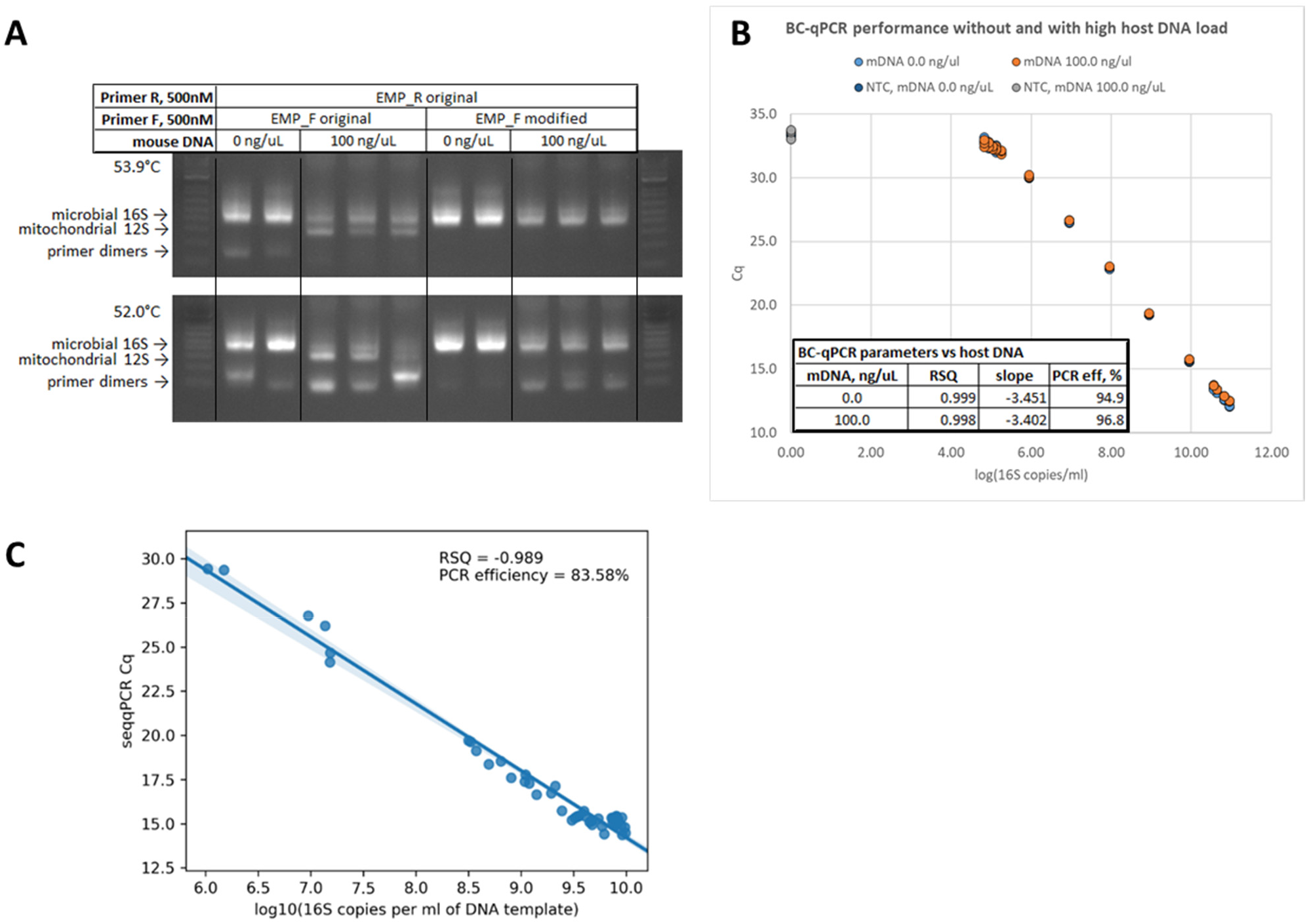
Optimization of the single-step protocol for microbial 16S rRNA gene DNA copy quantification and amplicon barcoding in samples without and with high mammalian DNA background. **(A)** Amplification products of the complex microbiota DNA sample containing 100 ng/µL of GF mouse DNA with the barcoded original EMP (UN00F0 + UN00R0) and barcoded modified (UN00F2 + UN00R0) primer sets. **(B)** Barcoding quantitative PCR reaction performance with the serial 10-fold dilutions of the complex microbiota DNA sample (SPF mouse fecal microbiota) with and without 100 ng/µL of mouse DNA assay. **(C)** Correlation of the BC-qPCR Cq values (Y-axis) with the absolute 16S rRNA gene DNA copy numbers (X-axis) previously determined in the same set of samples with and without high host DNA background (data in panel C are taken from [6, 7]) using the UN00F2 + UN00R0 qPCR assay.

The BC-qPCR assay demonstrated good performance in samples with and without high host DNA background (GF mouse DNA spiked in at 100 ng/µL of the DNA template sample) and containing the specific complex microbiota template (SPF mouse fecal DNA) across multiple orders of concentration (Fig. 4B). Regardless of the presence of high host DNA background, the reaction efficiency was ~95.0% and the assay was able to resolve 1.25 to 1.67-fold differences in total 16S rRNA gene copy loads among samples within the range of ~10^4.83^-10^10.95^ copies/mL.

We next confirmed that the BC-qPCR assay can provide accurate quantification data for the amount of 16S rRNA gene DNA copy loads in the analyzed samples. The Cq values obtained based on the real-time fluorescence measurements during the BC-qPCR reaction were in a good agreement with the absolute 16S rRNA gene DNA copy values (Fig. 4C) estimated in the same samples (samples and data were from [6, 7]) using the previously optimized qPCR assay (Fig. 3C).

### Single-step approach enables detection of absolute fold differences for quantitative microbiota profiling in lumenal and mucosal samples

The single-step BC-qPCR approach was used to calculate the absolute fold differences (as in Fig. 2C) for a number of taxa (Fig. 5) among samples from four experimental groups of mice described in [6, 7] using the NGS data from [6, 7] and the absolute fold difference data for the total 16S rRNA gene abundance in the corresponding samples from BC-qPCR assay (this study).

**Fig. 5.**
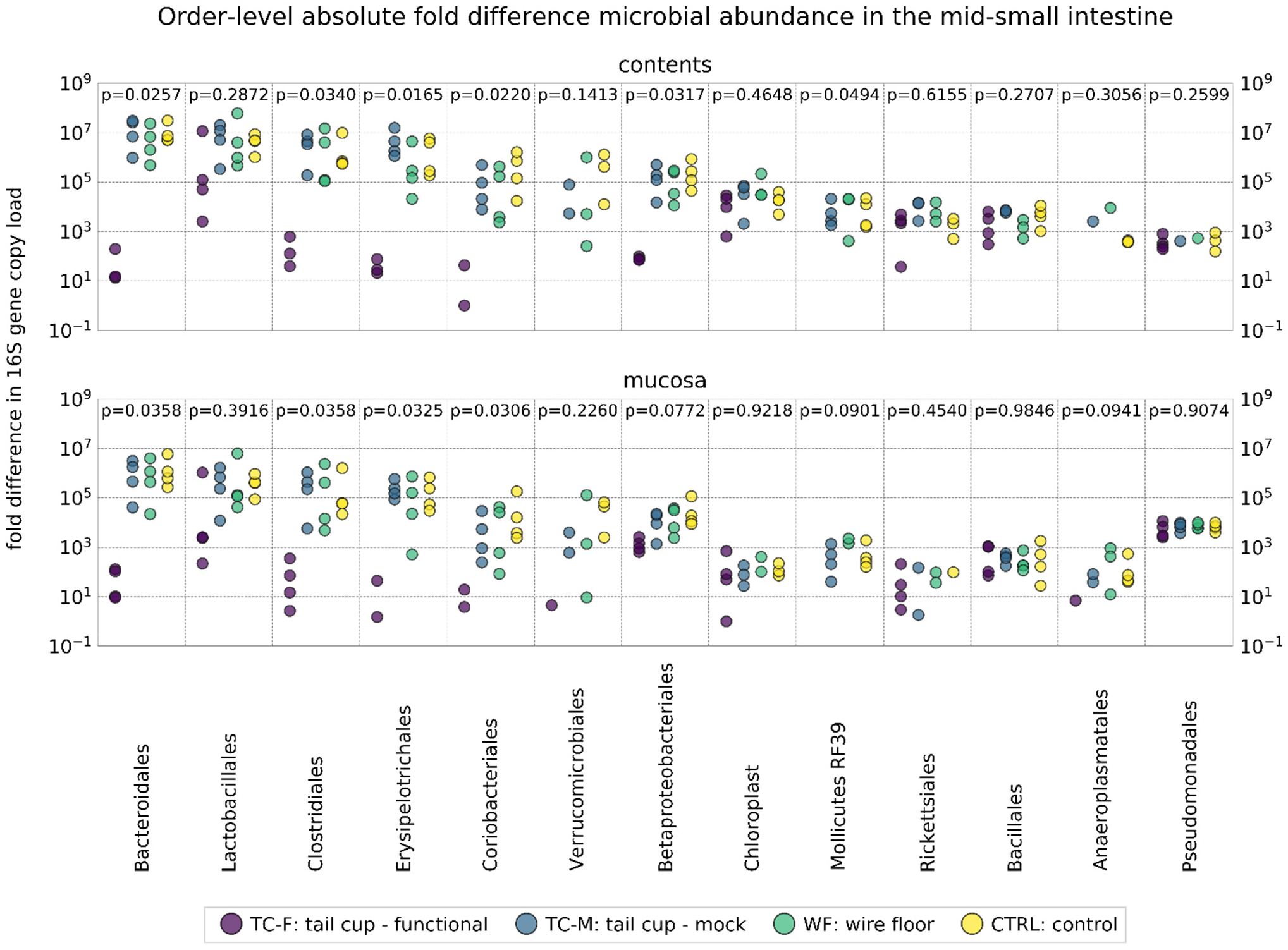
Absolute fold differences in the abundance of taxa (16S rRNA gene copies) in mouse mid-small intestine mucosal and lumenal samples yielded by the BC-qPCR assay. NGS data obtained from [6, 7] were used to calculate the fold difference values among samples using the single-step fold-difference approach (this study) for each individual taxon (order level). Multiple comparisons between the four experimental groups of mice were performed for each taxon using the Kruskal–Wallis test.

The absolute fold difference data for each individual taxon that are yielded by the single step approach can be used for comparisons among groups subjected to different experimental conditions using non-parametric rank tests (e.g., Kruskal-Wallis). Such comparisons revealed that in both the mid-small-intestine contents and mucosal samples multiple taxa (e.g., *Bacteroidales*, *Clostridiales*, *Erysipelotrichales*, *Coriobacteriales*) were differentially abundant (on the absolute scale) among the four experimental groups of mice (Fig. 5). These results are in agreement with data previously obtained using a two-step approach (utilizing absolute 16S rRNA gene copy quantification with a dedicated qPCR assay performed separately from the barcoding PCR reaction) described in [6, 7].

## DISCUSSION

Utilizing the uncharacterized samples for quantification anchoring (using the companion dPCR assay) and BC-qPCR efficiency estimation may be beneficial in order to account for the specific sample matrix effects or inhibitor carryover on the assay. However, if any dramatic sample-specific effects on the assay efficiency are observed (e.g., reaction inhibition) it is likely that not only the quantitative aspect of the BC-qPCR assay may be compromised, but the uniformity of the amplicon barcoding reaction may be significantly affected (e.g., introduced biases) as well. Thus, in order to take a full advantage of this platform it is important to ensure high quality and purity (e.g., from PCR inhibitors) of extracted DNA samples.

It is noteworthy that high host (e.g., mouse) DNA background per se does not inhibit the BC-qPCR assay even at very high concentrations (up to 100 ng/μL of template sample or DNA extract) as shown in Fig. 4B. Our work demonstrates that an uncharacterized extracted DNA sample with the total DNA concentration of up 100 ng/µL regardless of whether this is mostly host DNA with low amount of microbial template (e.g., mucosal biopsy) or mostly microbial DNA sample with some amount of host DNA (e.g., small-intestinal contents) can be reliably analyzed using the BC-qPCR assay.

## CONCLUSIONS

The single-step BC-qPCR approach enables accurate quantification of the number of 16S rRNA DNA gene copies and unbiased absolute abundance profiling of the microbial community structure in samples with microbial loads varying across multiple orders of magnitude and containing high host DNA background. The BC-qPCR approach offers the following advantages over the methods currently used in the field:

- Broader coverage of microbial diversity (87% bacteria, 87% of archaea based on the 16S rRNA marker gene sequences [4, 5]) maximizes the completeness of microbial detection and quantification and richness of diversity profiling.
- Microbial 16S rRNA gene DNA copy quantification demonstrated a broad dynamic range: the lower limit of quantification (LLOQ) – ~10^4.83^ copies/mL and the upper limit of quantification (ULOQ) – ~10^10.95^ copies/mL.
- Total 16S rRNA gene copy quantification has high resolution – ~1.25-1.67-fold differences in absolute 16S rRNA gene DNA copy concentrations can be distinguished in the demonstrated dynamic range with and without high host DNA background (100 ng/μL).
- “What’s quantifiable – is sequenceable, what’s sequenceable – is quantifiable”: this method maximizes correspondence between the total 16S rRNA gene DNA copy quantification data and 16S rRNA gene amplicon profiling data as a major advantage over the currently implemented approaches [8–12].
- Primer design allows for a good 16S rRNA gene DNA real-time (quantitative) PCR, digital PCR, and amplicon barcoding PCR reaction performance in samples with high mammalian host DNA background. No host DNA depletion is required for accurate microbial quantification and profiling, which is an advantage over the methods currently implemented in the field.
- Optimized single-step 16S rRNA gene DNA amplicon barcoding and quantification approach (performed in a single PCR reaction instead of two separate PCR reactions for quantification and barcoding) reduces the reagent and time costs while providing richer absolute or fold-difference microbiota profiles of the analyzed samples. Importantly, for samples with limited number of DNA copies, direct single-step quantification and barcoding eliminates the need for sample splitting, maximizing the utilization of precious samples.
- Optimized amplicon barcoding PCR reaction chemistry and workflow prevent amplification artifacts and biases [2] that could affect the accuracy of relative abundance measurements across samples with broad range of microbial loads and thus requiring different numbers of amplification cycles.
- The approach eliminates the need in synthetic spike-ins for accurate quantitative 16S rRNA gene amplicon profiling. Easily accessible commercial microbiome standards (e.g., ZymoBIOMICS) can be integrated as absolute 16S rRNA gene copy quantitative standards in the proposed method. Alternatively, uncharacterized samples analyzed with this approach can serve as absolute 16S rRNA gene copy quantification standards if a user is able to implement the companion dPCR assay described in this report.
- The approach may be applicable in both single (described in this report) and dual-indexing workflows.
- Overall, the proposed single-step method for quantitative 16S rRNA gene amplicon profiling based on the conventional real-time (quantitative) qPCR workflow allows for broad and immediate adoption of the approach in the field of basic and clinical microbiome research.

## METHODS

More detailed description of the methods is available in [6, 7].

### Animal samples

All animal handling and procedures were performed in accordance with the California Institute of Technology (Caltech) Institutional Animal Care and Use Committee (IACUC).

Fecal samples were collected from SPF C57BL6/J mice of 2-12 months of age originally purchased from Jackson Laboratory (Sacramento, CA, USA) and housed in the Caltech animal facility for up to 10 months. Germ free mouse intestinal mucosal samples from germ free C57BL6/J mice of 2-5 months of age obtained from the germ-free mouse colony maintained in the Caltech animal facility were collected and processed as in [6, 7].

All animal samples for Fig. 5 were obtained from [6, 7].

### DNA extraction

Total DNA was extracted from fecal and mucosal samples preserved in DNA/RNA Shield (DRS) solution (R1100-250, Zymo Research, Irvine, CA, USA) or fresh using ZymoBIOMICS DNA Miniprep Kit (D4300, Zymo Research) as described in [6, 7].

Quantitative (linear) recovery of DNA in the range of 16S rRNA gene copies of ~10^3.5^-10^11^ per ml was verified in using a series of 10-fold dilutions of specific-pathogen-free (SPF) mouse fecal microbial suspension in saline.

### Quantitative PCR (qPCR) for 16S rRNA gene DNA copy enumeration

This method is described in detail in [6, 7]. Briefly, each qPCR reaction was set up with 1.5 µL of DNA sample, qPCR master mix (SsoFast EvaGreen Supermix, #172-5200, Bio-Rad Laboratories, Hercules, CA, USA), forward (UN00F2, 5′-CAGCMGCCGCGGTAA-3′) and reverse (UN00R0, 5′-GGACTACHVGGGTWTCTAAT-3′ [1, 3]) primers (Integrated DNA Technologies, San Diego, CA, USA) at the final concentration of 500 nM each, and ultrapure water (Invitrogen UltraPure DNase/RNase-Free Distilled Water 10977-015, Thermo Fisher Scientific) to the final volume of 15 µL.

The thermocycling program was set up as follows: initial denaturation at 95 °C for 5 min. followed by 40 cycles each consisting of denaturation at 95 °C for 15 sec., annealing at 53 °C for 10 sec., and extension at 68 °C for 45 sec.

Assay was performed on a real-time PCR instrument (CFX96 Real-Time PCR Detection System, Bio-Rad Laboratories). The raw fluorescence data were processed and the Cq values were extracted with the accompanying software (Bio-Rad CFX Manager 3.1, #1845000, Bio-Rad Laboratories)

### Digital PCR (dPCR) for absolute 16S rRNA gene DNA copy enumeration

This method is described in detail in [6, 7]. Briefly, each reaction was set up with 2.0 µL of DNA sample, ddPCR master mix (QX200 ddPCR EvaGreen Supermix, #1864033, Bio-Rad Laboratories), forward (UN00F2, 5′-CAGCMGCCGCGGTAA-3′) and reverse (UN00R0, 5′-GGACTACHVGGGTWTCTAAT-3′ [1, 3]) primers (Integrated DNA Technologies) at the final concentration of 500 nM each, and ultrapure water (Thermo Fisher Scientific) to the final volume of 20 µL. In some experiments, additional DNA intercalating dye (EvaGreen, #31000, Biotium, Fremont, CA, USA) was added to the reactions up to ×1 final concentration (to achieve up to ×2 overall concentration). Each reaction volume was converted to droplets using a QX200 droplet generator (#1864002, Bio-Rad Laboratories).

Droplet samples were amplified on a thermocycler (C1000 Touch, #1841100, Bio-Rad Laboratories) according to the program: initial denaturation at 95 °C for 5 min. followed by 40 cycles each consisting of denaturation at 95

°C for 30 sec., annealing at 52 °C for 30 sec., and extension at 68 °C for 60 sec.; followed by the dye stabilization step consisting of 5 min incubation at 4 °C, 5 min incubation at 90 °C, and incubation at 12 °C for at least 5 min.

Droplet samples were quantified on a QX200 Droplet Digital PCR System (#1864001, Bio-Rad Laboratories) The raw data were analyzed and the target molecule concentrations were extracted using the accompanying software (QuantaSoft Software, #1864011, Bio-Rad Laboratories).

### 16S rRNA gene DNA amplicon barcoding for next generation sequencing (NGS)

This method is described in detail in [6, 7]. Briefly, all DNA samples were amplified and barcoded in triplicates for. Each reaction was set up with 3 µL of DNA sample combined with the PCR master mix (5PRIME HotMasterMix, #2200400, Quantabio, Beverly, MA, USA), ×1 DNA intercalating dye (EvaGreen, #31000, Biotium, Fremont, CA, USA), barcoded forward (UN00F2_BC, 5′-AATGATACGGCGACCACCGAGATCTACACTATGGTAATTGTCAGCMGCCGCGGTAA-3′) and reverse (UN00R0, 5′-CAAGCAGAAGACGGCATACGAGAT[NNNNNNNNNNNN]AGTCAGTCAGCCGGACTACHVGGGTWTCTAAT-3′ [1, 3], where [NNNNNNNNNNNN] – 12-nucleotide barcode sequences from [3]) primers (Integrated DNA Technologies) at the final concentration of 500 nM each, and ultrapure water (Thermo Fisher Scientific) to the final volume of 30 µL.

The thermocycling program was set up similarly to the EMP protocol [1, 3] as follows: initial denaturation at 94 °C for 3 min. followed by variable for each sample number of cycles each consisting of denaturation at 94 °C for 45 sec., annealing at 54 °C for 60 sec., extension at 72 °C for 105 sec.; followed by a final extension step at 72 °C for 10 min.

Assay was performed on a real-time instrument (CFX96 Real-Time PCR Detection System, Bio-Rad Laboratories).Samples were amplified for a variable number of cycles and each sample was removed from the heating block during the last 15 sec. of the current cycle extension step upon reaching the mid-exponential amplification phase. Each removed sample was maintained on a secondary heating block at 72 °C until all samples from the amplification series were amplified and returned to the primary heating block for the final extension step. The raw fluorescence data were processed and the fluorescence profiles over time were extracted with the accompanying software (Bio-Rad CFX Manager 3.1, #1845000, Bio-Rad Laboratories).

### Gel electrophoresis

Endpoint amplification products from whole PCR reactions were diluted 4-fold in ultrapure water (Invitrogen) and analyzed by gel electrophoresis using 1% (E-Gel EX, #G401001, Thermo Fisher Scientific) and 2% agarose gels (E-Gel, #G501802, Thermo Fisher Scientific).

### Barcoding PCR real-time data processing

Raw fluorescence data were processed using Python tools (Python tools used are described and referenced in [6, 7]).

Amplification profiles (fluorescence) for each PCR sample replicate were baseline-corrected by subtracting the minimal fluorescence value from the first 15 amplification cycles for each amplification replicate. Baseline-corrected amplification profiles from all replicates were averaged for each sample. Baseline-corrected and averaged amplification profiles were then used to find the Cq values (cycle numbers) at which they reached the fluorescence threshold (chosen as 2000 RFU) by interpolation.

Cq values were converted to absolute fold-difference values in total 16S rRNA gene copy load using the equations 3.1 and 3.2 (Fig.2) and assuming the BC-qPCR efficiency of 95.0%. The absolute fold difference values were then used to convert the taxa 16S rRNA gene relative abundance data obtained from the next generation sequencing to the taxa 16s rRNA gene absolute fold-difference data.

### Digital PCR (dPCR) for Illumina library quantification

This method is described in detail in [6, 7]. Briefly, a home-brew digital PCR library quantification assay was set up using the Illumina P5 and P7 adapter sequences as priming sites [1, 3, 13–15].

Each reaction was set up with 2.0 μL of the diluted amplicon sample ligated with the Illumina adapters, ddPCR master mix (QX200 ddPCR EvaGreen Supermix, #186-4033, Bio-Rad Laboratories), forward (ILM00F(P5), 5′-AATGATACGGCGACCACCGA-3′,) and reverse (ILM00R(P7), 5′-CAAGCAGAAGACGGCATACGA-3′,) primers (Integrated DNA Technologies) at the final concentration of 125 nM each, and ultrapure water (Invitrogen) to the final volume of 20 µL.

Thermocycling program was set up as follows: initial denaturation at 95 °C for 5 min, followed by 40 cycles each consisting of denaturation at 95 °C for 30 sec. and annealing-extension at 60 °C for 90 sec.; followed by the dye stabilization step consisting of 5 min incubation at 4 °C, 5 min incubation at 90 °C, and incubation at 12 °C for at least 5 min.

Hardware setup and droplet analysis were performed as in “Digital PCR (dPCR) for absolute 16S rRNA gene DNA copy enumeration”.

### Library pooling, purification, and quality control

This method is described in detail in [6, 7]. Briefly, triplicates of each barcoded amplicon sample were combined. After each sample was quantified with the home-brew ddPCR library quantification assay and KAPA SYBR FAST Universal qPCR Library Quantification Kit (#KK4824, Kapa Biosystems, Wilmington, MA, USA), all samples were pooled in equimolar quantities. The library was purified with Agencourt AMPure XP beads (#A63880, Beckman Coulter, Brea, CA, USA) and eluted with ultrapure water (Invitrogen). Quality control on the pooled library was performed using light absorbance at 260/280 nm (NanoDrop 2000c, Thermo Fisher Scientific) and the mean amplicon size of ~400 nucleotides was confirmed with a High Sensitivity D1000 ScreenTape System (#5067-5584 and #5067-5585, Agilent Technologies, Santa Clara, CA, USA) on a 2200 TapeStation instrument (Agilent Technologies) supported with the Agilent 2200 TapeStation Software A02.01. (Agilent Technologies).

### Next generation sequencing

This method is described in detail in [6, 7]. Briefly, paired-end 300-base reads were generated on a MiSeq instrument (Illumina, San Diego, CA, USA) using a MiSeq Reagent Kit v3 (#MS-102-3003, Illumina) with a PhiX control spiked in at 15%.

The following sequencing primers were used:

- MiSeq read 1: Seq_UN00F2_Read_1, 5′-TATGGTAATTGTCAGCMGCCGCGGTAA-3′.
- MiSeq read 2: Seq_UN00F2_Read_1, 5′-AGTCAGTCAGCCGGACTACHVGGGTWTCTAAT-3′ [1, 3].
- MiSeq index read: Seq_UN00R0_RC_Index, 5′-ATTAGAWACCCBDGTAGTCCGGCTGACTGACT-3′ [1, 3].

### Sequencing read processing and sequencing data processing

All processed and analyzed NGS data were obtained from [6, 7].

## Data Availability

NGS data and sequencing data analysis scripts were obtained and are available from [6, 7].

## Declaration of Interests

The contents of this article are the subject of a patent application filed by Caltech.

## Acknowledgements

This work was supported in part by the Kenneth Rainin Foundation Innovator Award (2018-1207), Army Research Office (ARO) Multidisciplinary University Research Initiative (MURI) contract #W911NF-17-1-0402, and the Jacobs Institute for Molecular Engineering for Medicine. We thank Natasha Shelby for contributions to writing and editing this manuscript.

