## Supplemental Figure S1 for "Quantitative microbiome profiling in lumenal and tissue samples with broad coverage and dynamic range via a single-step 16S rRNA gene DNA copy quantification and amplicon barcoding"

**Supplementary Figure S1**

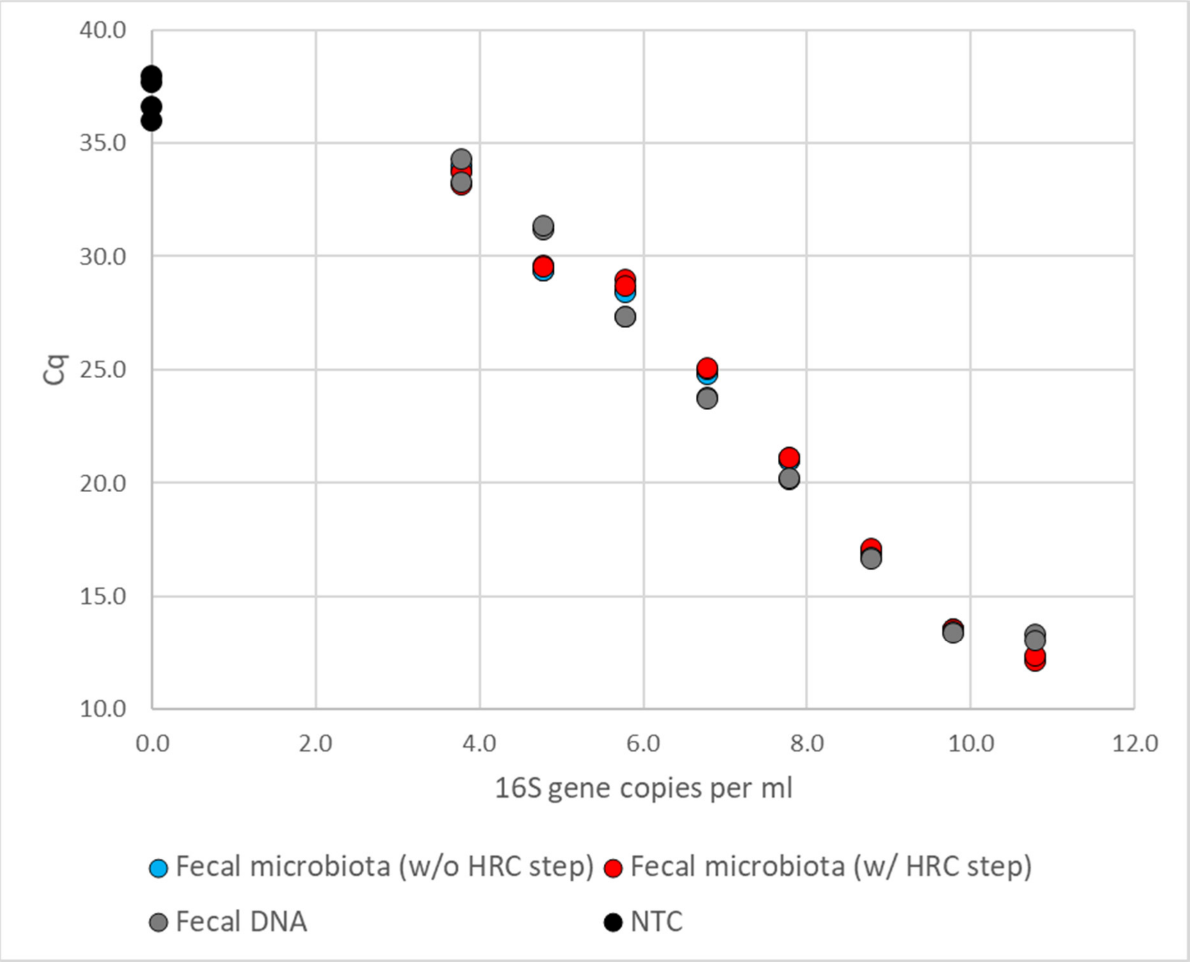


**Fig. S1. Quantitative DNA recovery using commercial extraction and purification kit (ZymoBIOMICS) from samples containing fecal microbial cells in the range of concentrations evaluated with a qPCR assay**. "Fecal microbiota (w/o HRC step)" – serial 10-fold dilutions of mouse fecal microbial suspension extracted with the kit with the HRC purification step omitted (N=1 extraction per dilution, N=3 PCR replicates). "Fecal microbiota (w/ HRC step)" – serial 10-fold dilutions of mouse fecal microbial suspension extracted with the kit and purified from PCR inhibitors using the "HRC" columns included with the kit (N=1 extraction per dilution, N=3 PCR replicates). "Fecal DNA" – serial dilutions of the single extracted DNA sample from the undiluted mouse fecal microbial suspension extracted with the kit according to the manufacturer's protocol (N=1 sample per dilution, N=3 PCR replicates). "NTC" – no-template control (N=4 PCR replicates).
